# Communication between DNA and nucleotide binding sites facilitates stepping by the RecBCD helicase

**DOI:** 10.1101/190215

**Authors:** Vera Gaydar, Rani Zananiri, Or Dvir, Ariel Kaplan, Arnon Henn

## Abstract

Double-strand DNA breaks are the severest type of genomic damage, requiring rapid response to ensure survival. RecBCD helicase in prokaryotes initiates processive and rapid DNA unzipping essential for break repair. Yet, the energetics of RecBCD during translocation along the DNA track needs to be quantitatively clarified. Specifically, it’s essential to understand how RecBCD switches between its binding states to enable its translocation. Here we determine, by systematic affinity measurements, the degree of coupling between DNA and nucleotide binding to RecBCD. We show that, in the presence of ADP, RecBCD binds weakly to DNA that harbors a double overhang mimicking an unwinding intermediate. Consistently, RecBCD binds weakly to ADP in the presence of the same DNA. We did not observe coupling between DNA and nucleotide binding for DNA molecules having only a single overhang, suggesting that RecBCD subunits must both bind DNA to “sense” the nucleotide state. Excitingly, we observed weak coupling for AMPpNp as RecBCD remains strongly bound to DNA in its presence. Detailed thermodynamic analysis of RecBCD reaction mechanism suggests an ‘energetic compensation’ between RecB and RecD, which may be essential for rapid unwinding. Our findings provide the basis for a ‘stepping mechanism’ during the processive translocation of RecBCD.

## Introduction

Double Strand DNA Breaks (DSDBs) are the severest type of genome damage, requiring fast and efficient repair (1). Helicases play an essential role in the repair mechanisms of DSDBs in every living organism (2). In prokaryotes, members of the RecBCD family unwind DSBs in preparation for strand invasion, which is essential for repair by homologous recombination (3). RecBCD is a highly processive DNA helicase exhibiting an exceptionally high unwinding rate of ∼ 1,600 base pairs (bp) per second (s^-1^) (4). RecBCD is a heterotrimer composed of one copy of RecB, RecC, and RecD, out of which RecB and RecD are DNA translocases and helicases (5–7). The RecC subunit “staples” the RecB and RecD subunits (8) and is crucial in destabilizing the duplex DNA ahead of the translocase activities of RecB and RecD (9) and recognizing the regulatory Chi sequence (10). RecBCD catalyzes ssDNA translocation on opposite DNA polarities, with RecB moving on the 3’ → 5’ strand and RecD moving on the 5’ → 3’strand, resulting in net translocation of RecBCD complex along the duplex DNA (6, 11, 12). RecBCD can also push through a highly crowded protein environment while maintaining processivity providing an additional enzymatic adaption to its mechanism(13). Although repair mechanisms for DSBs are present in all living organisms, there is no known eukaryotic homolog of RecBCD in terms of its structural organization or rapid and processive unwinding.

For RecBCD to transverse processively with the two motors translocating along the individual strands of the DNA, a stepping-like mechanism is most likely required. For example, a simple alternation between strong and weak DNA binding states can support processive translocation(13–15). The continuous threading and sliding of ssDNA through the proposed DNA tunnels or cavities in RecBCD will most likely advance via alteration between attached (strong binding state) and detached (weak binding) states. In this sense, RecBCD could function as a double-headed ATPase molecular machine, i.e., a molecular motor that has two motors that transverse along the DNA lattice, in analogy to double-headed molecular motors such as myosin V, which has been shown to perform processive stepping along the actin filament by walking “hand over hand” (16–18). Threading ssDNA through a semi-open DNA tunnel that forms in RecB and partially in RecD will increase processivity but still requires cycles of coordinated association-dissociation transitions to permit net vectorial translocation (19). The bound nucleotide state (ATP, ADP-P_i_, or ADP) may support such a mechanism by modulating the motor affinities towards the lattice it traverses on, as shown for all myosins and kinesin, as for some helicases (20–26). Establishing the existence of nucleotide intermediates during the mechanochemical ATPase cycle, and quantifying the free energy associated with them, is essential to understand the processive and rapid translocation of RecBCD.

In this work, we systematically determined the affinity of RecBCD for DNA substrates in the presence and absence of nucleotides. We characterized RecBCD’s binding to diverse DNA substrates and nucleotides, mimicking intermediate states along its unwinding reaction cycle to reveal the nucleotide-binding linkage within RecBCD. In most states, RecBCD complexes exhibit weak coupling between the nucleotide and the DNA bound to the RecBCD complexes. However, we found a strong coupling between the binding of ADP and dohDNA (a DNA construct mimicking an unwinding intermediate). Significantly, we observed a ∼40-fold reduction in affinity for RecBCD•ADP towards dohDNA. In the reciprocal experiment, RecBCD•dohDNA exhibits a much weaker affinity towards ADP, consistently showing the same strong coupling. Our results suggest that an ADP-bound biochemical intermediate, where strong coupling between DNA and ADP binding is observed, is essential to allow stepping during the translocation of RecBCD.

## Materials and Methods

### Reagents, expression, and purification of RecBCD

All chemicals and reagents were the highest purity commercially available. ATP and ADP were purchased from Roche Molecular Biochemicals (Indianapolis, IN, USA). Adenosine 5′-(β,γ-imido)triphosphate (AMPpNp) was purchased from Sigma (St. Louis, MO, USA). A molar equivalent of MgCl_2_ was added to nucleotides immediately before use. Nucleotide concentrations were determined by absorbance using an extinction coefficient e_259_ of 15,400 M^-1^ cm^-1^. The concentrations of N-methylanthraniloyl (mant) derivatives of ADP, 2’-deoxyADP, ATP, and 2’-deoxyATP (Jena Bioscience, Jena, Germany) were determined using e_255_ of 23,300 M^-1^ cm^-1^. Unless otherwise specified, all experiments were conducted in RecBCD Buffer (RB: 20 mM MOPS pH 7.4, 75 mM NaCl, 2 mM MgCl_2_, 1 mM DTT). Over-expression and purification of recombinant RecBCD was based on the previously described method (23, 24), with an additional step described by Zananiri *et al.* (27). The RecBCD concentration was determined using e_280_ = 4.2 ×10^5^ M^-1^ cm^-1^ in guanidinium chloride. RecBCD purity of nucleic acid contaminants was determined by measuring the absorption ratio of 280/260 nm, and only protein fractions with a ratio >1.3 were used.

### DNA substrates preparations

DNA oligonucleotides were purchased from IDT (Leuven, Belgium) and were HPLC purified. The DNA substrates are shown in Fig S1. All DNA used in the binding experiments was obtained by folding or hybridizing the DNA in 20 mM MOPS pH 7.4, 75 mM NaCl, and 1 mM MgCl_2_ buffer at 85° C for 3 minutes, followed by slow cooling to room temperature before storage at -20° C.

### DNA binding measurements by Fluorescence Anisotropy (FA)

FA measurements were performed with a PC1 spectrofluorometer set up T-format configuration for simultaneous acquisition on two emission channels using monochromators equipped with automatic polarizers. Samples were equilibrated in RB for 60 min at RT and then measured with α_ex_ = 492 nm using vertically polarized light. The emitted vertical and horizontal polarized light was monitored at 90° with emission monochromators at α;_em_ =523 nm at 25 ± 0.1°C. The instrument manufacturer calculated the g-factor to correct the gains between vertical and horizontal PMT detectors. Nucleotide concentrations in the indicated measurements were 2 mM (Mg•ADP, or Mg•AMPpNp). Fluorescent DNA substrates were held at a constant concentration of 25 nM.

### RecBCD oligomeric state and stoichiometric binding isotherms

To rule out the existence of a dimer of RecBCD during catalysis, we determined the oligomeric state of RecBCD under our experimental conditions, similar to what had been performed earlier (10). Fig. S2A shows the stoichiometric binding of ssDNA and hpDNA (Fig. S1 & Table S1) under the condition in which [DNA] >> *K*_D_ and RecBCD is titrated well above the *K*_D_ (Table 1) measured by FA (Fig. 1A) and fitted according to Eq. 1:

**Figure 1:**
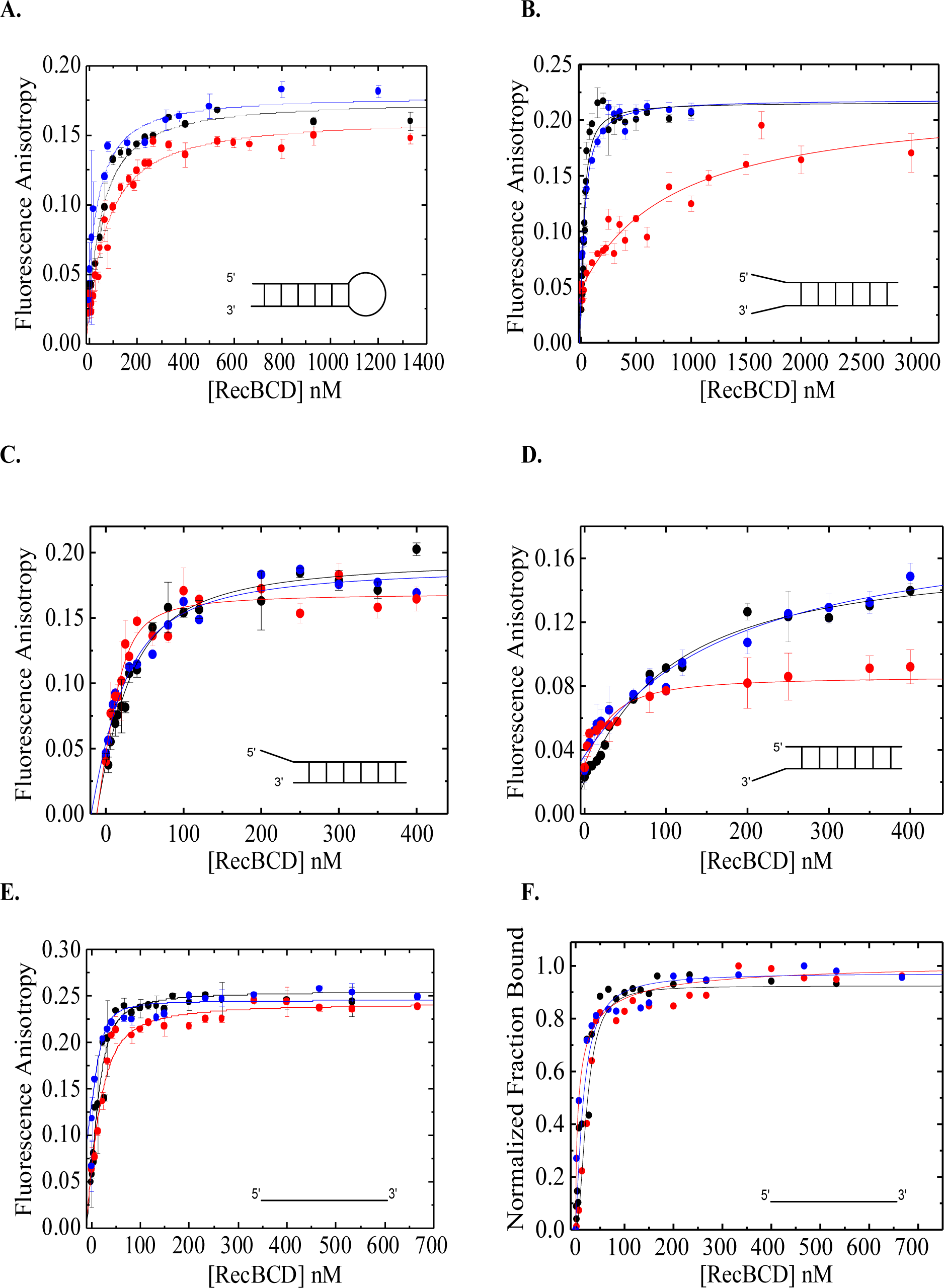
Fluorescence Anisotropy equilibrium binding measurements of RecBCD and RecBCD×nucleotides complex to fluorescently labeled DNA. DNA substrates were labeled with 6-fluorescein amidite. Fluroescent DNA was held at 25 nM and RecBCD was titrated to reach saturation of the binding isotherm. Binding to RecBCD (•), RecBCD×AMPpNp (•), and RecBCD×ADP (•) is shown for A. hpDNA, B. dohDNA, C. 5ohDNA, D. 3ohDNA and E. ssDNA. λ_ex_ = 492 nm and λ_em_ = 523 nm. Nucleotides concentrations where indicated were 2 mM (Mg×ADP, or Mg×AMPpNp). F. Data in B fitted to a Hill equation. Error bars report the s.d. of n = 3 independent measurements. Solid lines show the best fit to Eq. 2.

**Table 1:**
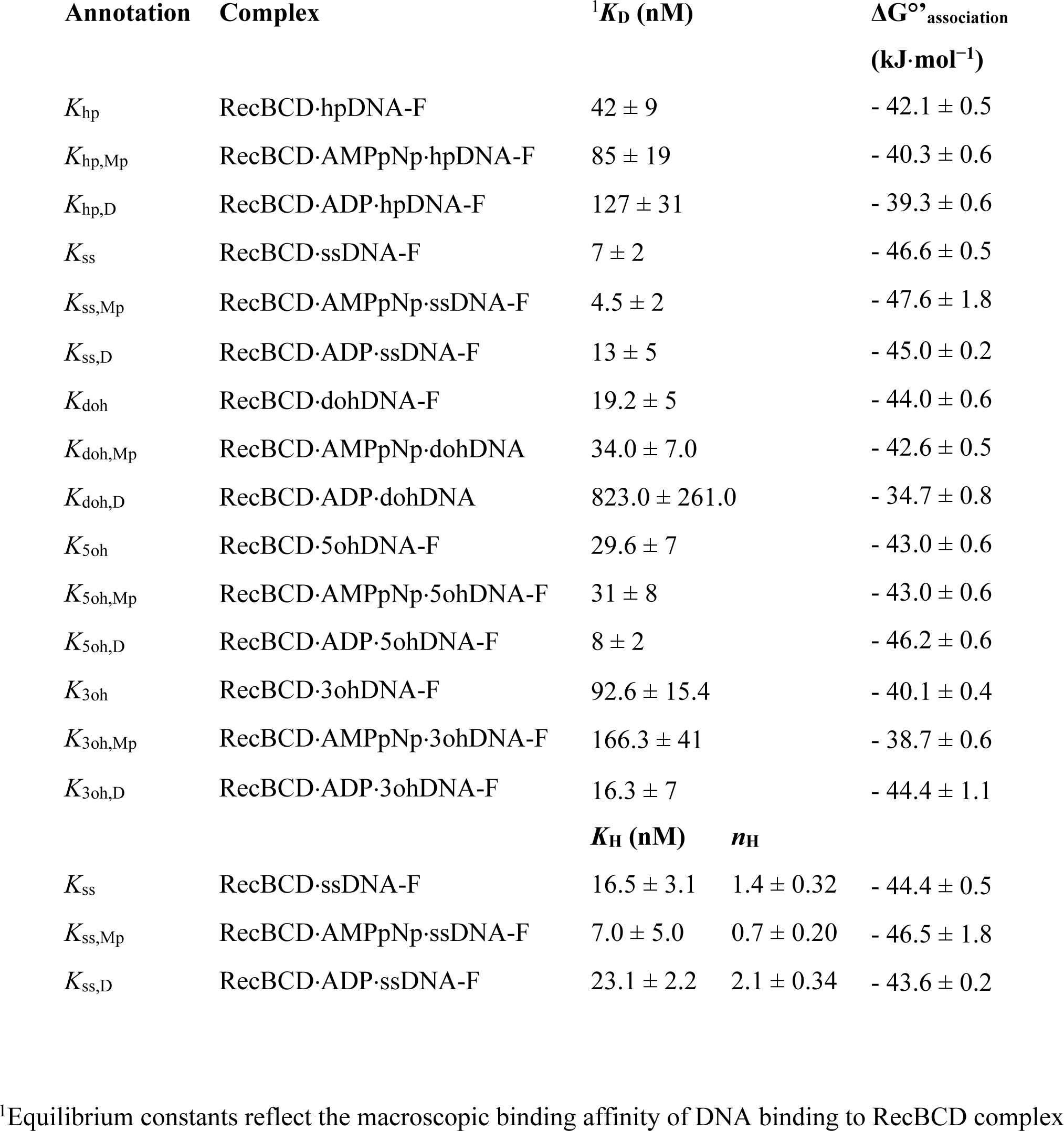
Equilibrium constants for DNA substrates binding to RecBCD and RecBCD×nucleotides complex.

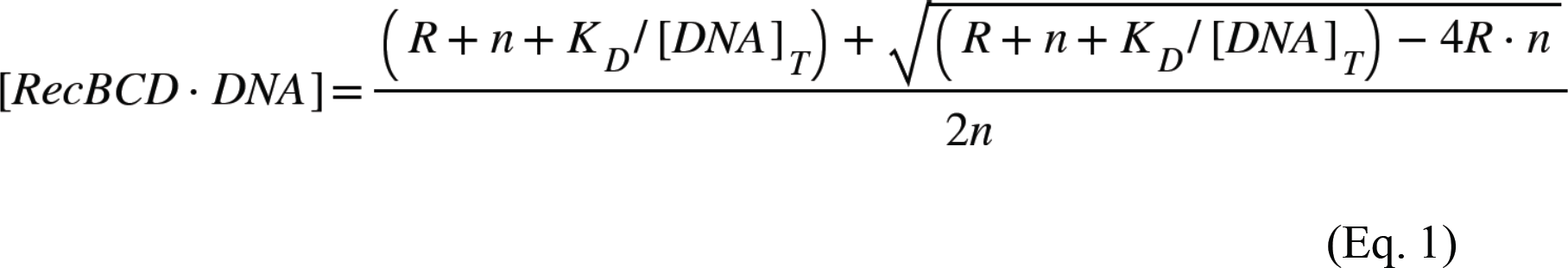

Under stoichiometric conditions of [DNA]_T_>> K_D_, R is the plotted (experimental) mole/mole ratio of RecBCD/DNA, n is the ratio of the fit of RecBCD to DNA, and the DNA_T_ is the DNA (ssDNA or hpDNA) concentration. This analysis shows that RecBCD:hpDNA and RecBCD: ssDNA binds with 1:1 and 1:2 stoichiometric ratio. Previously, the stoichiometry binding of fluorescently labeled single-stranded overhang (Fig. S1C) had a similar stoichiometry ratio to the RecBCD heterotrimer (28).

### Analysis by analytic size exclusion chromatography (SEC)

RecBCD•hpDNA complex in comparison to RecBCD (SI Fig. 2B) clearly shows a monodispersed peak of approximate MW of ∼330 kDa (determined by the linear range of known MW protein marker) in complex with the major peak of the hpDNA with a ratio of 1:1, 280:260 nm absorbance (in comparison to only RecBCD of 3:1, 280:260 ratio). This further confirms the RecBCD: hpDNA stoichiometry to be 1:1.

**Figure 2:**
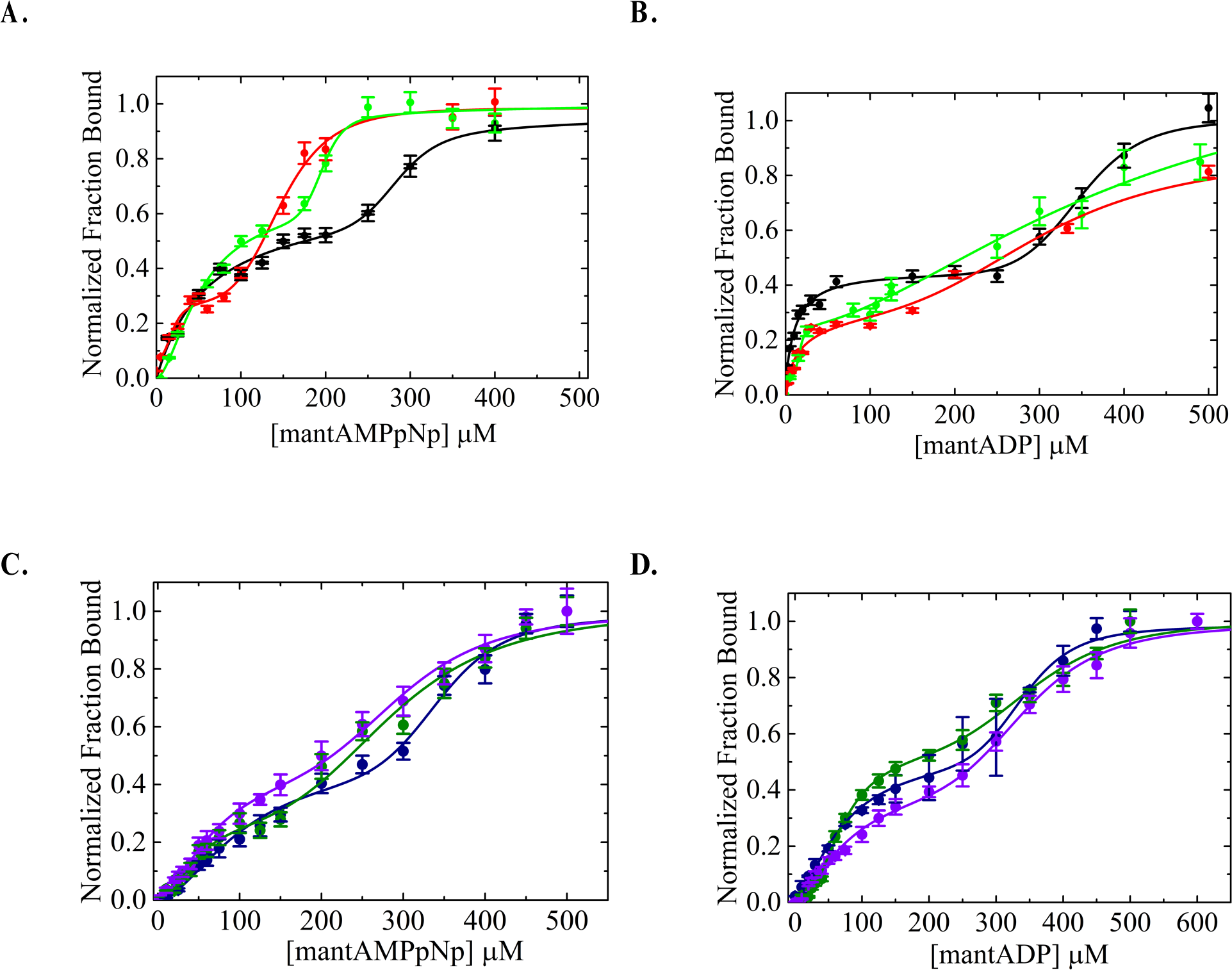
Equilibrium binding of RecBCD and RecBCD×DNA to mantNucleotides. **A. and B.** Titration curves of mantAMPpNp (A) and mantADP (B) binding to RecBCD (•), RecBCD×hpDNA (•), and RecBCD×ssDNA (). The data for RecBCD×mantAMPpNp (A) and RecBCD×mantADP (B), collected contemporaneously to the rest of the data, is adapted from (27). **C. and D.** Titration curves of mantAMPpNp (C) and mantADP (D) binding to RecBCD×dohDNA (e), RecBCD×5ohDNA (e), and RecBCD×3ohDNA (e). Solid lines show the best fit to Eq. 3. Error bars report the s.e.m. of n = 3 independent measurements.

### DNA binding isotherms for *K*_D_ determination

#### Model 1

The binding of DNA substrates to RecBCD is described by the following stoichiometric reaction scheme (Scheme 1):

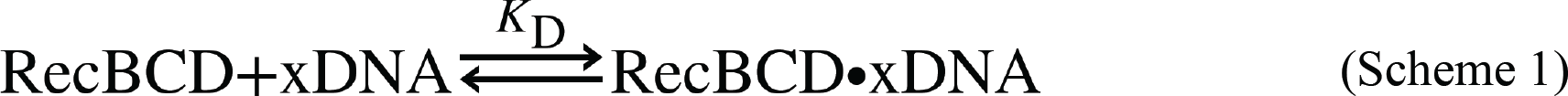

Where xDNA can be any of the DNA substrates used in this work The general solution for this equilibrium binding scheme is in the form of the following quadratic Eq. 2:

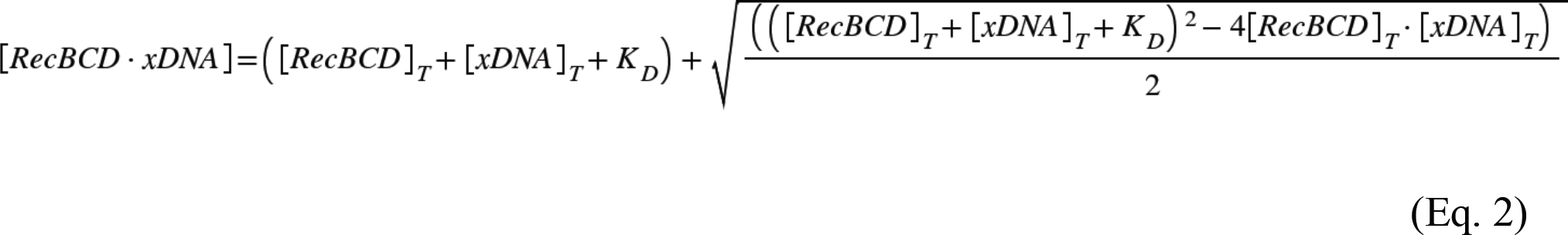

Where [DNA]_T_ is the monitored species; [RecBCD]_T_ is the titrated species, [RecBCD•xDNA] is the bound species. For FA measurements, the premise is that the total fluorescence intensity remains constant throughout the titration (27, 29, 30). Under our measurement conditions, the changes in the FTI were minimal to be neglected, allowing the direct fitting of the FA binding isotherm curve using Eq. 2 to determine the equilibrium binding constants.

#### Model 2

The analysis of ssDNA binding to RecBCD with the Hill equation reveals cooperativity in the presence of ADP. RecBCD binds two ssDNA. Therefore, we can utilize a binding model such as Hill to account for multiple binding sites according to Scheme 2:

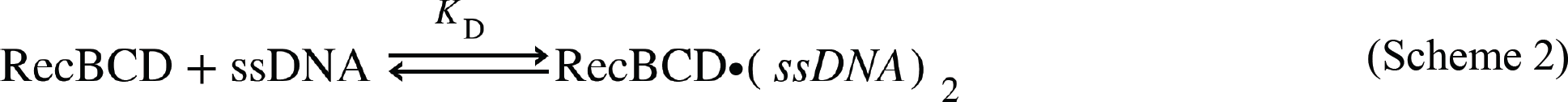

To extract the two parameters *K*_H_ and *n*_H_ as shown in Equation 3:

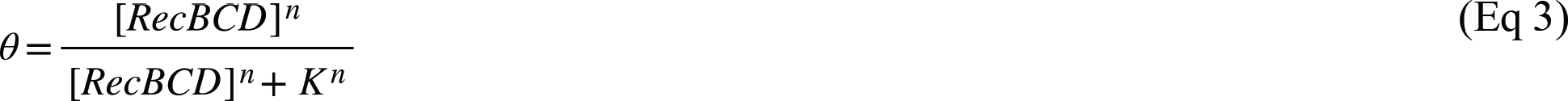

Where θ is the fraction bound *K* is the apparent dissociation constant, and *n* is the Hill coefficient, assuming that the ssDNA binding sites are similar but not identical.

### mantNucleotide binding to RecBCD by Förster resonance energy transfer (FRET)

FRET measurements were performed using a PC1 spectrofluorometer (ISS, Champaign, IL) with excitation and emission monochromators. The observation cell was regulated with a Peltier temperature controller at 25 ± 0.1°C. All equilibrium binding reactions were performed in a 10 μl Precision cell fluorescence cuvette (Farmingdale, NY, USA), which allows minimal inner filter effects up to a concentration of ∼ 550 mM mantNucleotides. The buffer included 20 mM MOPS pH 7.4, 2 mM MgCl_2_, 1 mM DTT, and varying concentrations of NaCl (75, 150, 200, 300 mM) (27). mantNucleotides were made with an equal concentration of MgCl_2_ (1:1 stoichiometric ratio) before being added to RB. Equilibrium binding reactions of mantNucleotides to RecBCD were measured by FRET between RecBCD intrinsic tryptophan fluorescence (._ex_ = 280 nm) and bound mantNucleotide (fluorescence monitored at 90° through an emission monochromator at λ_em_ = 436 nm). We subtracted the background fluorescence of free nucleotides on the observed emission peak. DNA substrates were constant at 1 mM, and RecBCD concertation was 1 mM. In the case of ADP binding to RecBCD•dohDNA, the DNA concentration was 8 mM. mantNucleotide binding curves of the fluorescence change as a function of the free ligand concentration are best fitted by the sum of two Hill plots according to Equation 4 (27):

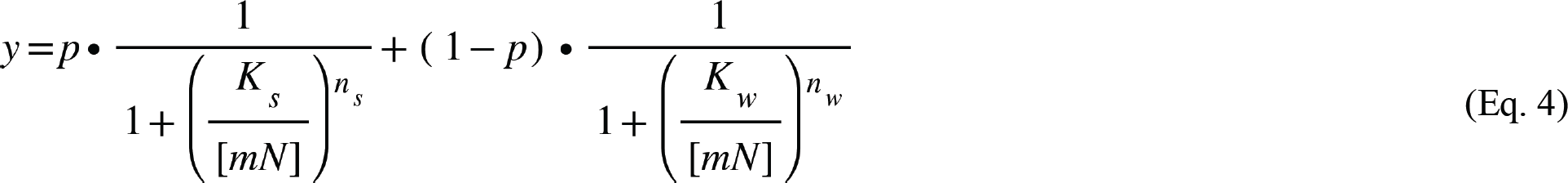

Where y is the fraction bound, *mN* is the ligand concentration, *K*_s_ (strong nucleotide states), and *K*_w_ (weak nucleotide states) are the equilibrium constants of the first and second phases, respectively. *n_s_* and *n_w_* are the Hill coefficients of the first and second phases, respectively, and *p* is the proportionality constant (0 ý *p* ý 1). Previously, we showed that the first phase of the binding titration curve reflects binding to the canonical ATP binding sites residing within RecB and RecD. The second phase describes binding to additional weak binding sites. Both are given by macroscopic equilibrium constant’s *K*_s_ and *K*_w_ (27).

## Results

### DNA binding affinities to RecBCD and RecBCD•nucleotide complexes

We used Fluorescence Anisotropy (FA) to measure the equilibrium binding affinity of RecBCD for DNA in the absence and presence of AMPpNp and ADP. To study various points in the catalytic unwinding reaction cycle, we devised a set of DNA substrates that mimic major biochemical states: binding of RecBCD to a DNA hairpin with a blunt end (hpDNA, Figure S1A) simulates the initiation phase; binding to a DNA hairpin with two non-complementary overhangs (dohDNA, Figure S1B) represents the DNA fork separation and unwinding; In addition, two substrates with either 5’ or 3’ overhangs (5ohDNA and 3ohDNA, respectively, Figure S1C, D) are used to measure the affinity of RecBCD for DNA when only RecB or RecD subunits can access and bind the ssDNA (8, 28, 31) (8, 24, 27), and a single-stranded DNA substrate (ssDNA, Figure S1E) is used to mimic the translocating complex. All the DNA binding isotherms exhibited hyperbolic dependence curves (Figure 1), enabling us to determine the equilibrium binding constants for the different DNA substrates (Table 1).

### ADP modulates the affinity of RecBCD for DNA

The binding of RecBCD to hpDNA, which mimics the initiation step of the catalytic cycle, revealed that the presence and identity of a nucleotide cofactor modulate the complex affinity. Specifically, the ADP bound state showed a ∼3-fold weaker affinity for hpDNA than the APO one, while an intermediate affinity was observed in the presence of AMPpNp (Fig. 1, Table 1). The strong affinity of the APO state agrees with earlier results showing the ability of RecBCD to bind and melt blunt-ended duplex DNA in an Mg^2+^-dependent, ATP-independent manner (31). Moreover, the reduction in affinity for the AMPpNp and ADP states observed here may provide a mechanism for RecBCD to translocate, following ATP binding and hydrolysis, from an initial relatively strong binding state to a DSDB site (32).

When we tested the binding of RecBCD to dohDNA, we observed a substantial reduction in the affinity in the presence of ADP (40-fold and 25-fold) compared to the APO and AMPpNp states, respectively; Fig. 1B and Table 1). This strong coupling between ADP and dohDNA binding indicates that RecBCD cannot bind strongly simultaneously to both ADP and dohDNA, i.e., the binding of one weakens the affinity of the other and vice versa.

To further dissect the basis for the observed weak binding to dohDNA in the presence of ADP, we characterized two additional DNA substrates containing either a 5ohDNA or 3ohDNA, likely engaging only with RecD or RecB, respectively (Fig. S1D, E). Generally, RecBCD exhibited a significantly higher affinity towards the asymmetric constructs than for dohDNA (Fig. 1B-D and Table 1). Specifically, RecBCD retained a strong affinity towards 5ohDNA and 3ohDNA in the presence of AMPpNp, and RecBCD•ADP displayed an even higher affinity for these DNA constructs. This suggests a weak coupling between the asymmetric substrates and the nucleotide state. We note here that the lower FA absolute value observed for the 3ohDNA in the presence of ADP in Figure 1D may be due to a less rigid conformation of the fluorophore at the 3’ overhang than a fluorophore at the 5’ overhang.

The observed differences in the binding affinities of the Apo state of RecBCD towards ssDNA, dohDNA, 5ohDNA, and 3ohDNA suggest that these four complexes interact specifically with their cognate binding sites matching the polarity and orientation of the DNA strands with RecB and RecD polarity (31–33). Hence, RecBCD engages with either 5’ or 3’ overhang DNA strands with a small effect by ADP binding. However, when both overhang DNA are engaged simultaneously with RecBCD (dohDNA substrate), ADP binding strongly weakens RecBCD affinity towards DNA, suggesting communication between RecBCD subunits. On the other hand, in the presence of AMPpNp (Fig. 1B-D, Table 1), a less pronounced effect is observed on the binding isotherms between the different substrates. Rather than weakening the binding affinities observed towards dohDNA in the presence of ADP, here, the overall affinity towards dohDNA is approximately the average of the individual binding affinities (34.0 ± 7.0, 31 ± 8, & 166.3 ± 41 nM for RecBCD•AMPpNp*X* dohDNA/5ohDNA/3ohDNA, respectively).

### Interaction of RecBCD with ssDNA revealed cooperative binding

We analyzed the binding isotherms of ssDNA to RecBCD with a hyperbolic binding curve assuming no cooperativity (Figure 1E, Eq.2). Treating the binding of both ssDNAs with one macroscopic constant showed that none of the nucleotide cofactors affected the affinity of RecBCD towards ssDNA binding (Figure 1E, Table 1). However, since ssDNA binds to RecBCD with a 2:1 stoichiometric ratio (Fig. S2), we can also use the Hill equation to fit our binding isotherms (Figure 1F, Table 1). ssDNA binding to RecBCD without nucleotides shows intermediate cooperativity and a Hill coefficient of *n*_H_ ∼ 1.4, whereas, in the presence of AMPpNp, the cooperativity is slightly negative (*n*_H_ ∼ 0.7). Interestingly, while ADP binding weakens the affinity towards ssDNA, it increases the Hill coefficient to *n*_H_ ∼ 2. Hence, when the two DNA binding sites in RecB and RecD are bound to ssDNA, the ADP nucleotide cofactor imposes a different degree of cooperativity, suggesting a different conformation and degree of interaction between RecB and RecD upon the different nucleotide cofactors. This may result from disrupting long-range intramolecular signaling between the three subunits as a response to different nucleotides (34).

### Nucleotide-binding to RecBCD and RecBCD•DNA complexes exhibit different degrees of affinities

Having characterized how nucleotides modulate the affinity of RecBCD for different DNA substrates, we turned to characterize how engaging with DNA modulates the affinity of RecBCD for nucleotides. We used FRET between mantNucleotide derivatives and intrinsic tryptophan residues and measured equilibrium binding curves (27). We previously showed that the binding of mantNucleotide to RecBCD and RecBCD•DNA complexes displays a biphasic binding isotherm curve to multiple binding sites (27). These macroscopic binding constants describe the two types of nucleotide binding sites: high affinity binding to the catalytic sites (in RecB and RecD) and low affinity binding to “auxiliary” sites that increase the flux of ATP towards the catalytic sites.

When we measured isotherms of ADP and AMPpNp binding to the different RecBCD-DNA complexes, the high-affinity phase displayed a binding constant, *K*_s,_ ranging from ∼13 to 106 mM and a Hill coefficient *n*_s_ between 0.8 and 3.0 (Table 2). The second weak binding phase constant, *K*_w,_ was twice as weak, exhibiting values of ∼140 to 350 mM (Table 2). In addition, these sites showed a degree of cooperativity, *n*_w_ ranging from 2.5 to 12.0, with significant errors, suggesting that there may be some cooperative binding (Table 2). We identified four RecBCD•DNA biochemical intermediates that differentially affected nucleotide binding, producing RecBCD•DNA•nucleotide complexes with different affinities (Table 2).

**Table 2:** Equilibrium constants for nucleotides binding to RecBCD and RecBCD×DNA complexes.

| Annotation | Complex | $K_s (\mu\text{M})$ | $n_s$ | $K_w (\mu\text{M})$ | $n_w$ | $p$ | $\Delta G_s^{\circ\prime}$<br>(kJ·mol<sup>-1</sup>) | $\Delta G_w^{\circ\prime}$<br>(kJ·mol<sup>-1</sup>) |
| --- | --- | --- | --- | --- | --- | --- | --- | --- |
| $^4K_{\text{Mp}}$ | RecBCD·mMp | $52.4 \pm 7.8$ | $0.99 \pm 0.1$ | $286.9 \pm 11.9$ | $12.0 \pm 4.7$ | $0.60 \pm 0.04$ | $-24.4 \pm 0.4$ | $-20.2 \pm 0.1$ |
| $K_{\text{Mp,hp}}$ | RecBCD·hpDNA·mMp | $73.5 \pm 9.3$ | $1.44 \pm 0.3$ | $147.0 \pm 6.8$ | $5.0 \pm 1.1$ | $0.74 \pm 0.1$ | $-23.6 \pm 0.3$ | $-21.9 \pm 0.1$ |
| $K_{\text{Mp,ss}}$ | RecBCD·ssDNA·mMp | $20.0 \pm 5.9$ | $2.90 \pm 0.9$ | $146.9 \pm 6.8$ | $5.0 \pm 1.1$ | $0.23 \pm 0.1$ | $-26.8 \pm 0.7$ | $-21.9 \pm 0.1$ |
| $K_{\text{Mp,doh}}$ | RecBCD·dohDNA·mMp | $106.0 \pm 33.5$ | $1.8 \pm 0.4$ | $343.8 \pm 9.2$ | $9.0 \pm 3.2$ | $0.50 \pm 0.1$ | $-22.7 \pm 0.8$ | $-19.6 \pm 0.2$ |
| $K_{\text{Mp,5oh}}$ | RecBCD·5ohDNA·mMp | $47.8 \pm 22.1$ | $2.0 \pm 1.0$ | $285.1 \pm 17.9$ | $4.1 \pm 0.8$ | $0.30 \pm 0.1$ | $-24.6 \pm 1.1$ | $-19.8 \pm 0.1$ |
| $K_{\text{Mp,3oh}}$ | RecBCD·3ohDNA·mMp | $71.0 \pm 28.3$ | $1.7 \pm 0.4$ | $295.8 \pm 12.1$ | $5.2 \pm 1.5$ | $0.50 \pm 0.1$ | $-23.7 \pm 1.0$ | $-20.2 \pm 0.2$ |
| $^4K_{\text{mD}}$ | RecBCD·mD | $13.2 \pm 3.1$ | $0.95 \pm 0.1$ | $322.5 \pm 6.9$ | $6.2 \pm 0.6$ | $0.45 \pm 0.03$ | $-27.8 \pm 0.6$ | $-20.1 \pm 0.1$ |
| $K_{\text{D,hp}}$ | RecBCD·hpDNA·mD | $29.2 \pm 11.8$ | $0.81 \pm 0.2$ | $314.9 \pm 22.0$ | $4.0 \pm 0.3$ | $0.51 \pm 0.12$ | $-25.9 \pm 1.0$ | $-20.0 \pm 0.2$ |
| $K_{\text{mD,ss}}$ | RecBCD·ssDNA·mD | $15.7 \pm 3$ | $2.85 \pm 1.7$ | $343.0 \pm 84.2$ | $2.5 \pm 0.8$ | $0.19 \pm 0.01$ | $-27.4 \pm 0.5$ | $-19.8 \pm 0.6$ |
| $K_{\text{mD,doh}}$ | RecBCD·dohDNA·mD | $70.9 \pm 19.4$ | $1.5 \pm 0.3$ | $338.3 \pm 8.9$ | $9.0 \pm 3.0$ | $0.51 \pm 0.1$ | $-23.7 \pm 0.7$ | $-19.8 \pm 0.1$ |
| $K_{\text{mD,5oh}}$ | RecBCD·5ohDNA·mD | $70.5 \pm 8.6$ | $2.4 \pm 0.4$ | $354.4 \pm 17.7$ | $5.3 \pm 1.1$ | $0.38 \pm .01$ | $-23.7 \pm 0.3$ | $-19.7 \pm 0.1$ |
| $K_{\text{mD,3oh}}$ | RecBCD·3ohDNA·mD | $80.8 \pm 25.8$ | $1.5 \pm 0.3$ | $342.2 \pm 7.5$ | $5.9 \pm 1.2$ | $0.44 \pm 0.2$ | $-23.3 \pm 0.8$ | $-19.8 \pm 0.1$ |

1. mantADP (mD); mantAMP-pNp (mMp);
2. Fitted parameters for the sum of two Hills equations, $K_s$ & $K_w$ are the equilibrium dissociation constants for nucleotides association to the strong and weak binding sites, respectively; $n_s$ & $n_w$ are the Hill coefficients for nucleotides' association to the strong and the weak binding sites, respectively.
3. $\Delta G_s^{\circ\prime} = RT \ln(K_s)$ and $\Delta G_w^{\circ\prime} = -RT \ln(K_w)$ are the standard binding energies at 1 M ligand, R is the gas constant (8.314 J mol<sup>-1</sup> K<sup>-1</sup>), T = 298.15°K for the strong and weak binding sites, respectively.
4. Previously reported (27).

AMPpNp binds to the strong binding sites in the RecBCD•dohDNA complex with two-fold weaker affinity than RecBCD (*K*_Mp,doh_ =106 ± 33.5 vs. *K*_Mp_ =52.4 ± 7.8, respectively, Fig. 2, Table 2). In contrast, the affinities towards RecBCD•5ohDNA and RecBCD•3ohDNA (*K*_Mp,5oh_ = 47.8 ± 22.1 and *K*_Mp,3oh_ 71.0 ± 28.3 respectively, Fig. 2A-D, Table 2) are similar to the one for RecBCD. These observations further suggest that the coupling between the DNA and nucleotide binding sites is more pronounced when both DNA binding sites are occupied.

Two major consequences are revealed by the binding of ADP towards RecBCD•DNA complexes. First, the overall affinity towards ADP is weakened in all substrates examined as compared to RecBCD (*K*_mD,doh_ = 70.9 ± 19.4, *K*_mD,5oh_ = 70.5 ± 8.6, *K*_mD,3oh_ = 80.8 ± 25.8 and *K*_mD_ = 13.2 ±3.1, respectively, Fig. 2, Table 2). Next, when the DNA substrates are physically connected ahead of the binding sites, there seems to be a more substantial impact on the nucleotide-binding affinities towards the strong binding sites. Therefore, it may well be that tension in the closed DNA form substrates remotely from RecBCD•DNA sites has a more substantial effect on intersubunit communication. Finally, ADP binding to RecBCD•DNA complexes are more affected than AMpNp binding to RecBCD•DNA (strong nucleotide binding sites affinities). In contrast to the modulation of nucleotide binding to the catalytic sites by engaging DNA, the affinity for the auxiliary sites shows no significant change between the RecBCD•DNA states of RecBCD and in the absence of any DNA substrates (Table 2). These results suggest that the weaker nucleotide binding sites (auxiliary sites) are uncoupled to nucleotide binding to the strong binding sites and are unaffected by DNA binding.

### Thermodynamic coupling constants between the DNA and nucleotide binding states reveal binding linkage within the RecBCD•DNA•nucleotide complex

The free energy derived from the binding constants of the RecBCD•DNA•nucleotide complex formation permits quantifying the degree of *linkage* and the thermodynamic coupling constant (TC) between any four states within a closed reaction cycle (35). Importantly, this analysis can provide the degree of coupling between the different states, primarily when a cofactor modulates the affinity towards a substrate or a substrate modulates the affinity towards the cofactor. In addition, it provides an estimation of the intrinsic consistency of a set of measurements, as the overall change in free energy ΔG° for a closed reaction cycle should be balanced, i.e., ΔG° = 0. Therefore, the TC equilibrium constant pairs within a balance thermodynamic coupling constants should be equal, i.e., *K*_Mp,hp_*/K*_Mp_ & *K*_hp,Mp_*/K*_hp_. Thus, ratios significantly > 1 report stronger coupling between the measured states, whereas ratios ∼ 1 report weak coupling, i.e., indicate that nucleotide binding does not affect the affinity of RecBCD towards DNA and *vice versa*. Figure S3 shows the ten closed reaction cycles corresponding to our five DNA substrates and two nucleotide states measurements. The results of TC analysis for the cycles of hpDNA, dohDNA, 5ohDNA, 3ohDNA, and ssDNA, in the absence or presence of either AMPpNp or ADP, are summarized in Table 3. The detailed thermodynamic squares are balanced for all the AMPpNp states and two states of ADP binding to RecBCD•hpDNA and RecBCD•ssDNA (Table 4 & Figure S3). AMPpNp seems to possess weak coupling with RecBCD; hence RecBCD can bind both the AMPpNp nucleotide and DNA quite strongly simultaneously. AMPpNp presumably mimics either the pre-hydrolysis bound ATP or the post-hydrolysis state ADP•Pi state, suggesting that RecBCD can remain strongly bound during these states to the DNA. However, ADP shows stronger coupling in the hpDNA and, to some extent, in the ssDNA. This suggests that these weaker binding states for ADP may be more dissociative biochemical intermediates when the ADP state is mainly populated during the ATPase cycle of RecBCD. Finally, for three thermodynamics squares: (*K*_D,doh_*/K*_D_)/(*K*_doh,D_*/K*_doh_), (*K*_D,5oh_*/K*_D_)/(*K*_5oh,D_*/K*_5oh_) and (*K*_D,3oh_*/K*_D_)/(*K*_3oh,D_*/K*_3oh_) (FigureS3, marked within yellow) the calculated ratios between the TC pairs are ∼0.13, ∼18, and ∼30, respectively. This suggests an unbalanced close reaction cycle with these three DNA substrates. Two factors may contribute to this: First, ADP binding is measured from the reverse pathway of the ATPase cycle, so it may not necessarily represent the same state after Pi’s release. In the kinetic scheme, ADP is bound when the ATPase cycle proceeds via a productive ATP hydrolysis forward pathway (36). Hence, a structural ADP state could not be accessible from the reverse binding of ADP to RecBCD. In addition, our model for nucleotide binding is based on mantNucleotides which may result in different affinity than unmodified nucleotides.

**Table 3:**
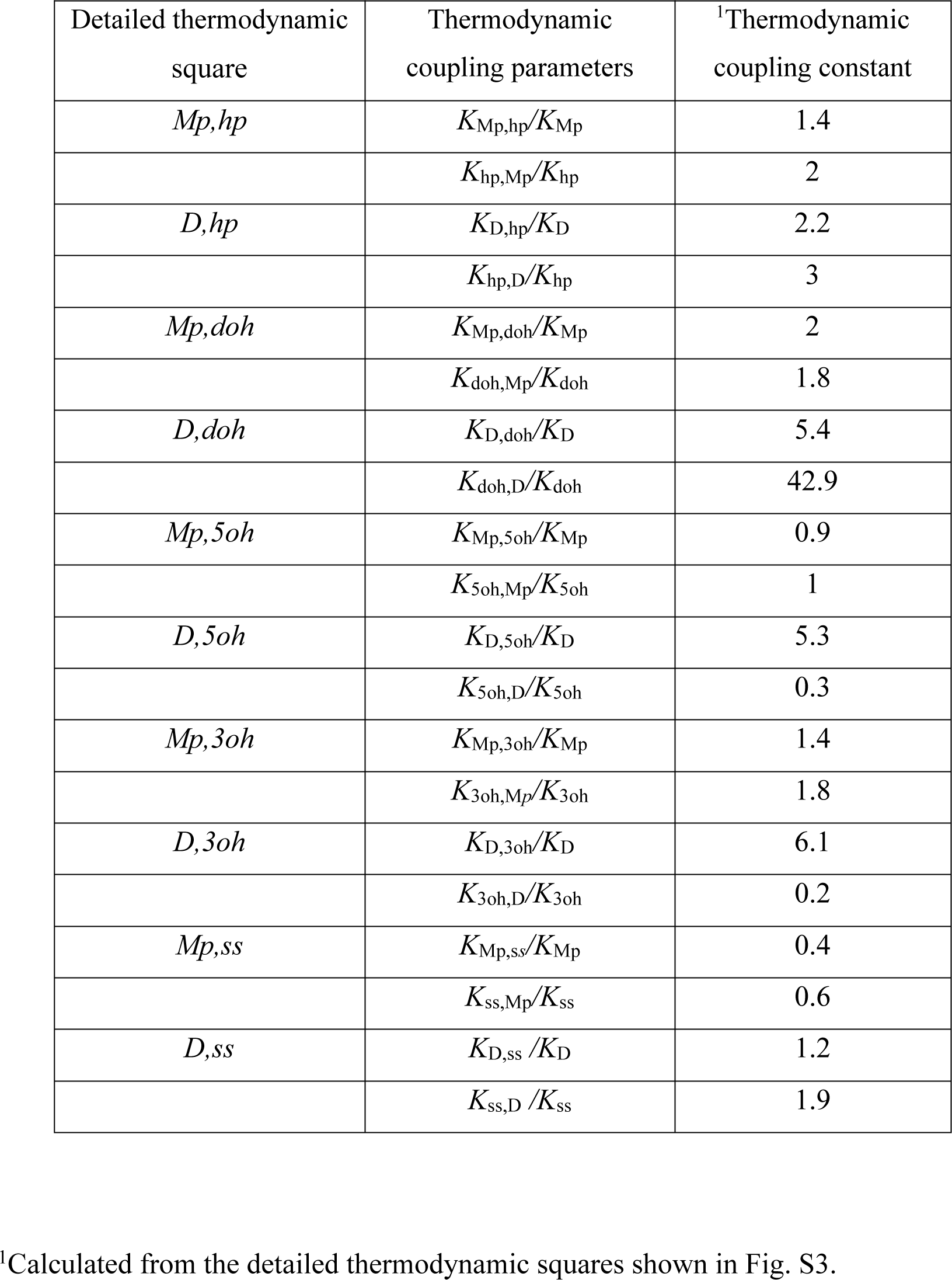
Thermodynamic coupling parameters associated with RecBCD, RecBCD×DNA, RecBCD×DNA× nucleotide complexes.

## Discussion

In this work, we have studied quantitatively the degree of coupling between DNA substrates mimicking unwinding intermediates and nucleotide states in RecBCD. Our results demonstrate that this coupling is significant enough to enable a stepping mechanism based on differential binding to the DNA lattice during highly processive and rapid translocation. ADP binding induces the strongest coupling compared to the APO or AMPpNp bound states. All three substrates, dohDNA, 5ohDNA, and 3ohDNA, affect ADP binding, unlike AMPpNp. When RecBCD is bound to ADP, we observed the most robust modulation in the binding affinity to all the substrates compared to the APO and AMPpNp states.

Our results also suggest that the occupancy of both RecB and RecD with bound ssDNA is essential for RecBCD intersubunit communication. Two significant observations support this requirement. First, the hpDNA substrate, which lacks ssDNA overhangs long enough to interact fully with RecB and RecD, does not show such an effect. Therefore, both ssDNAs beyond the fork must be strongly embedded within their binding sites to impose communication between nucleotide and DNA binding. The second observation stems from the binding affinity towards dohDNA, compared to that for either 5ohDNA or 3ohDNA. The 5ohDNA binds well in the presence of ADP; however, when ssDNA overhangs are present in both antiparallel strands, ADP has the opposite effect inducing weak binding towards dohDNA. The observation that single-stranded overhang substrates do not have the same impact on ADP binding to RecBCD implies that intersubunit communication between RecBCD subunits is induced to a fuller extent when both RecB and RecD are bound to ssDNA (8, 27, 32, 37).

Further evidence for the importance of occupancy of both RecB and RecD to propagate the observed TC is evident by the 3’ overhang DNA. In this case, the 3ohDNA substrate displays weaker affinity towards RecBCD in the absence or presence of AMPpNp than when ADP is bound. This suggests that AMPpNp may generate a weak binding state when one DNA binding site is occupied. Hence, ADP is not the sole cofactor to weaken DNA binding. However, the TC for these states, *K*_Mp,3oh_*/K*_Mp_, and *K*_3oh,Mp_*/K*_3oh_, is very weakly coupled, and the detailed balance is consistent with the thermodynamic cycle among these states. Corollary, there may be a weakening in the affinity among the abovementioned states, but they don’t trigger the strong coupling effects observed with ADP in the presence of dohDNA. Furthermore, when both subunits are engaged with ssDNA, the observed TC is most likely indicative of interacting residues residing remotely from the DNA binding sites within RecB and RecD (32).

Characterization of the major biochemical intermediates along the reaction coordinates of RecBCD allows us to formulate a minimal energetic model for the unwinding of a single ATP hydrolysis cycle (Figure 3). The calculated overall Δ Δ G° from RecBCD·dohDNA^N^ to RecBCD·dohDNA^N-1^ (i.e., the Δ Δ G° associated with unwinding and translocating 1 bp) is about ∼7 kJ•mol^−1^ which is about the energy released from the hydrolysis of one mole of ATP. Assuming that the ATPase cycles of RecB and RecD are unsynchronized and possess different nucleotide states during ATPase cycling, the overall Δ Δ G° would be an average of two different states at any given moment during the unwinding. This will minimize the transition barrier between states. For example, Pi release seems to be an unfavorable intermediate as the Δ Δ G° = +8.3 ± 1.0. However, ADP release is highly favorable Δ Δ G° =-9.3 ± 1.0. Hence, if, i.e., RecB releases Pi and RecD releases ADP, the net change is negative, ∼ Δ Δ Δ G° = -1.0, enabling RecBCD as complex to overcome unfavorable transition states. We proposed that this strategy by RecBCD will allow it to progress with minimum energy barriers as a holoenzyme.

**Figure 3:**
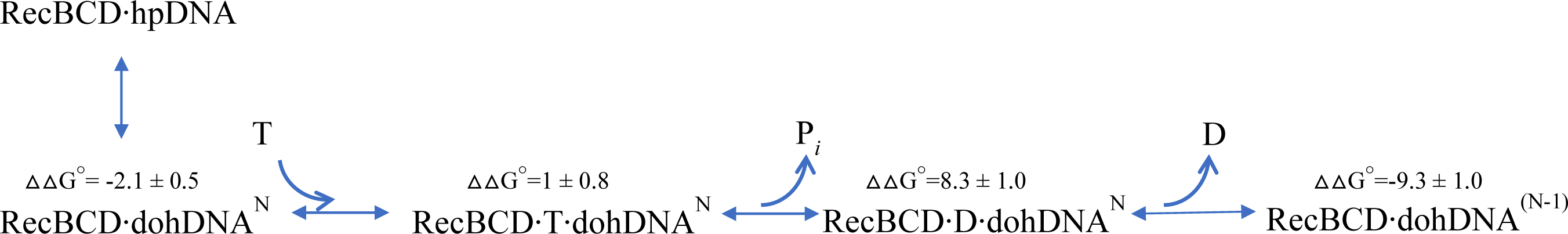
Biochemical intermediates along a discrete model for unwinding from state N (i.e. N bp unwound) to state N-1. The model assumes one ATP molecule is hydrolyzed per bp unwound. For each complex, ΔΔG° is calculated at saturating nucleotide concentration, from the change in ΔG°’_association−1_ of the two complex-forming reactions. All ΔΔG° are reported in kJ mol (Table 1).

Structural specifics of RecBCD DNA contact sites within the RecBCD subunits, capable of determining the identity of the residues involved in the observed TC, are still unavailable. In addition, there are still no structures of RecBCD with different nucleotide states and different substrates of DNA that can reveal the conformational changes induced by binding. However, inspecting the available RecBCD structure shows a large surface area directly interacting with the DNA (8, 32). The RecB “arm” at the forefront makes extensive contact with the fork created upon dsDNA binding and passive DNA melting (32). Additional contacts of the junction of the dsDNA with the ‘pin’ domain of RecC add to the overall RecBCD-DNA interaction. RecB 2A subdomain interacts with the 3’ overhang ssDNA, while RecD subdomains 1A and 2A interact with the 5’ overhang ssDNA. During the translocation of RecB and RecD motors in opposite directions, the contacts of both subunits with DNA dynamically change to allow movement. Such dynamics are driven by different nucleotide binding ligated states, modulating the affinity as resolved in this work. Strong binding states can be viewed as the ‘pulling’ states, while the weak binding states are the ‘relieving’ states allowing for productive advancement of the RecBCD motor, alternating these events between RecB and RecD. Notably, to result in processive translocation, the weak binding states (or strong binding states) cannot coexist in both subunits simultaneously. Hence allosteric communications likely exist and involve interactions across all the DNA contact sites throughout the three RecBCD subunits.

### Data availability

The authors confirm that the data supporting the findings of this study are available within the article and Supplementary Data.

## Supplementary data

Submitted separetly

## Acknowledgments

We thank Dr. Stephen C. Kowalczykowski and Dr. Theetha Pavankumar for the RecBCD expression system and their assistance in the RecBCD purification protocol (the University of California, Davis, California).

