## Supplementary Data for "Communication between DNA and nucleotide binding sites facilitates stepping by the RecBCD helicase"

**Title:**

### Supplementary Data.

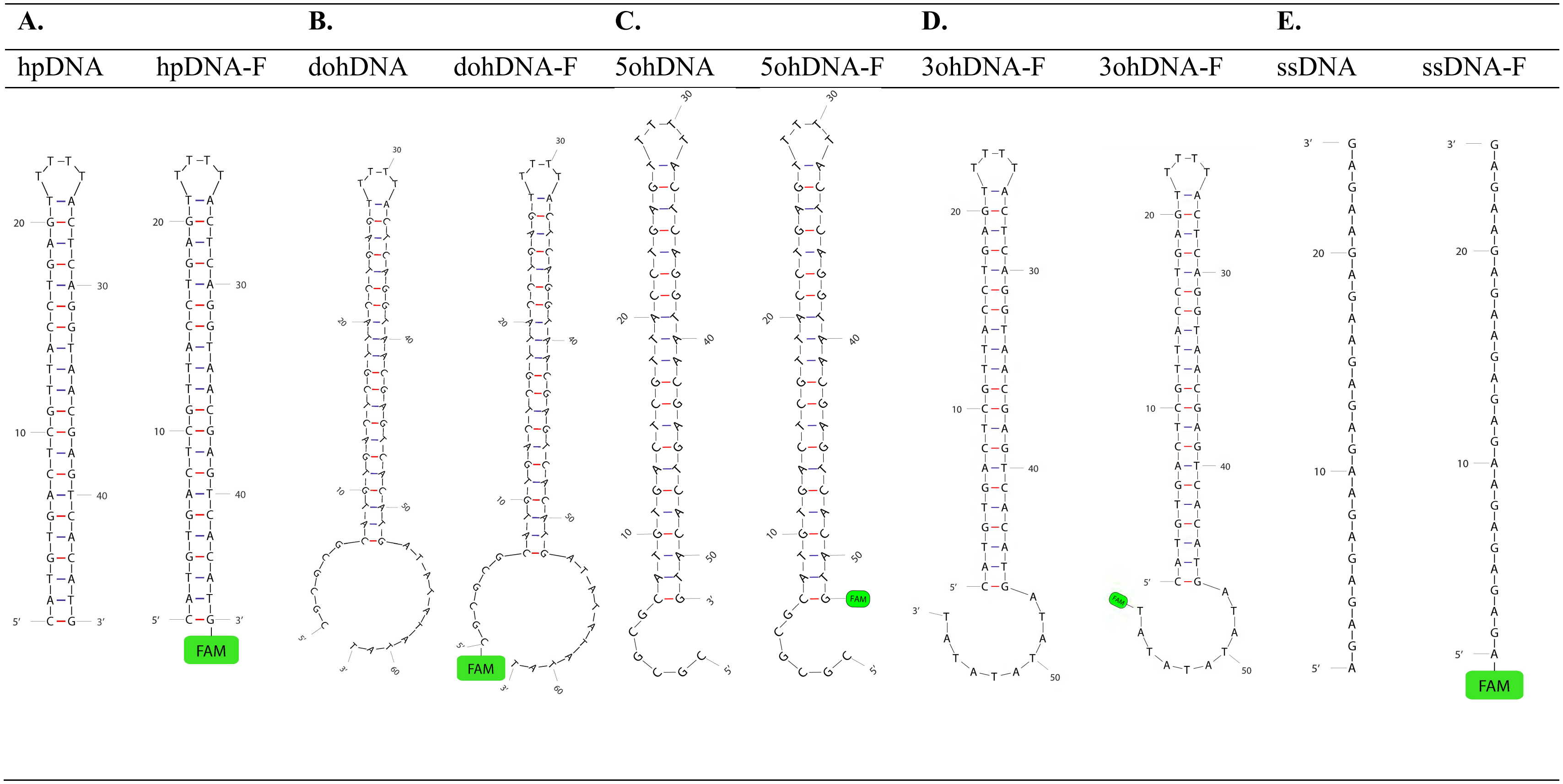


**Figure S1**: **Schematic representation of the DNA substrates used in this work**. **A.** Hairpin DNA: hpDNA, hpDNA-F (Table S1, 1a, b) **B.** Double strand overhang DNA: dohDNA, dohDNA-F (Table S1, 2a, b), **C.** Single strand 5'-overhang DNA: 5ohDNA, 5ohDNA-F (Table S1, 3a, b), **D.** Single strand 3'-overhang DNA: 3ohDNA, 3ohDNA-F (Table S1, 4a, b) **E.** Single strand DNA: ssDNA, ssDNA-F (Table 1, 5a, b). F=FAM(6-fluorescein amidite)

**A.** **B.**

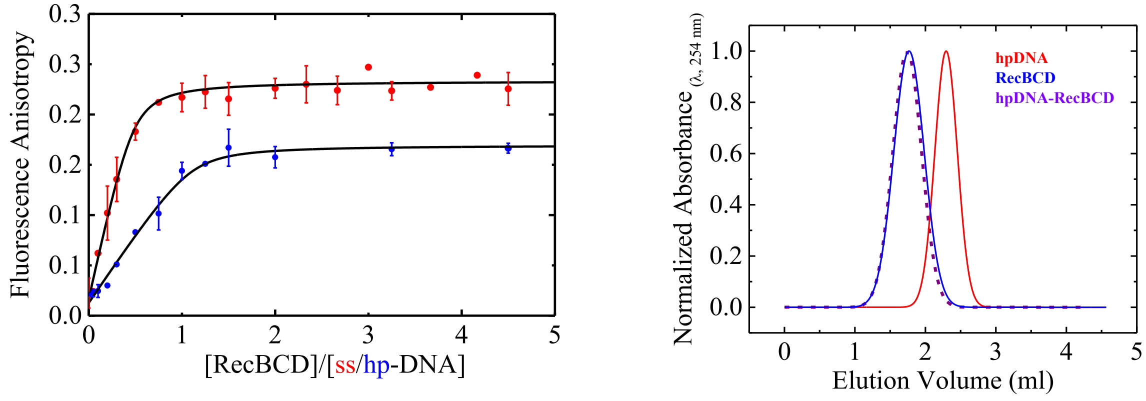


**Figure S2: Purification and Biochemical characterization of RecBCD oligomeric states without and with DNA and nucleotides. A.** Stoichiometric titration of RecBCD versus ssDNA and hpDNA using FA. Concentration of ssDNA/hpDNA were fixed at 2 μM, and RecBCD was titrated up to 10 μM (5-fold DNA concentration). The fitted curve shown in black line is according to Eq. 1. This produced the following "n" values for RecBCD/ssDNA = 0.49 ± 0.1 and RecBCD/hpDNA = 1.12 ± 0.1. Data shown as mean ± s.e.m., n=2. **B.** Analytical SEC analysis by Superdex 200 of RecBCD oligomeric state in the absence and presence of hpDNA under stoichiometric conditions. The absorption spectra were normalized for their presentation. RecBCD absorption ratio 280 nm:260 nm for RecBCD in the absence of hpDNA is 3:1, in the presence hpDNA the ratio is 1:1, therefore enable the determination of hpDNA presence in RecBCD peak. Peak of RecBCD·hpDNA, corresponding to a molecular weight of ~ 330 kDa according to the MW standards.

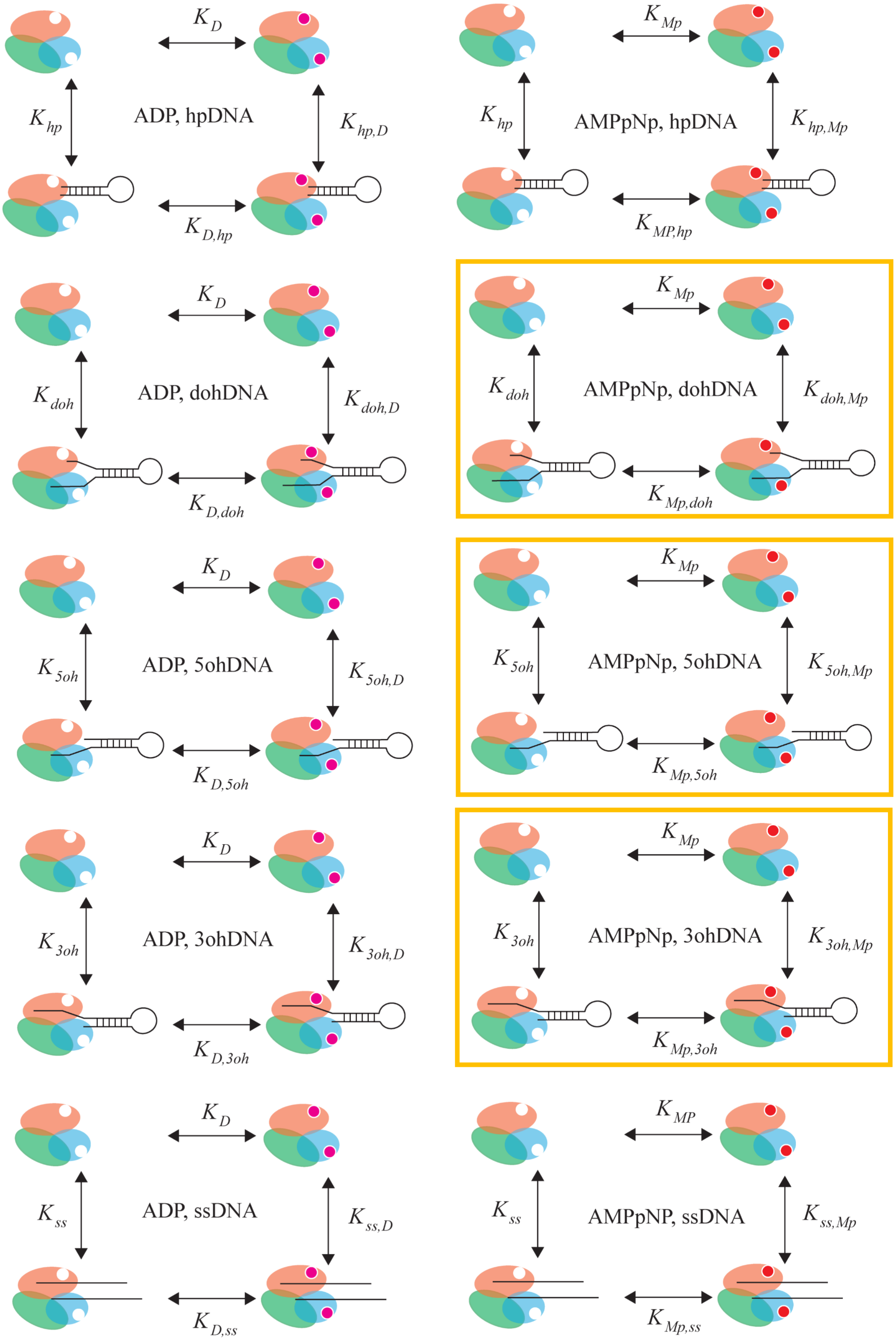

$K_{D}$
$K_{hp}$
ADP, hpDNA
$K_{hp,D}$
$K_{D,hp}$
$K_{Mp}$
$K_{hp}$
AMPpNp, hpDNA
$K_{hp,Mp}$
$K_{MP,hp}$
$K_{D}$
$K_{doh}$
ADP, dohDNA
$K_{doh,D}$
$K_{D,doh}$
$K_{Mp}$
$K_{doh}$
AMPpNp, dohDNA
$K_{doh,Mp}$
$K_{Mp,doh}$
$K_{D}$
$K_{5oh}$
ADP, 5ohDNA
$K_{5oh,D}$
$K_{D,5oh}$
$K_{Mp}$
$K_{5oh}$
AMPpNp, 5ohDNA
$K_{5oh,Mp}$
$K_{Mp,5oh}$
$K_{D}$
$K_{3oh}$
ADP, 3ohDNA
$K_{3oh,D}$
$K_{D,3oh}$
$K_{Mp}$
$K_{3oh}$
AMPpNp, 3ohDNA
$K_{3oh,Mp}$
$K_{Mp,3oh}$
$K_{D}$
$K_{ss}$
ADP, ssDNA
$K_{ss,D}$
$K_{D,ss}$
$K_{MP}$
$K_{ss}$
AMPpNP, ssDNA
$K_{ss,Mp}$
$K_{Mp,ss}$

**Figure S3: Detailed thermodynamic balance schemes of RecBCD ligated states.** The schemes describe the ligated states of the initiation complex (first row, hpDNA, stoichiometry 1:1), translocation complex (second row, ssDNA, stoichiometry 1:2), unwinding complex (third row, dohDNA, stoichiometry 1:1). The boxes in the fourth and fifth rows are asymmetric substrates engaged either with RecD (5ohDNA) or RecB (3ohDNA) only. The equilibrium constants, and hence the nucleotides binding occupancy in the model correspond only to the strong nucleotides binding sties. The equilibrium constants and the free energy calculated are shown in Tables 2 and 3 and are according to the detailed balanced shown in the reaction schemes. H – RecBCD, hp – hpDNA, ss – ssDNA, doh – dohDNA, 5oh – 5ohDNA, 3oh – 3ohDNA, D – ADP, Mp – AMPpNp. The RecBCD enzyme is represented by orange, green, blue ovals for RecB, RecC and RecD, respectively. DNA is drawn as black solid lines.

**Table S1: Sequences of the DNA substrates used for DNA and nucleotides binding experiments.**

| # | Sequence Description | Sequence 5'- 3' | DNA Length |
|---|---|---|---|
| 1a | 21 bp hairpin DNA | CATGTGACTCGTTACCTGAGTTTTTACTCAGGTAACGAGTCACATG | 46 |
| 1b | 21 bp hairpin DNA 3' FAM | GATGTGACTCGTTACCTGAGTTTTTACTCAGGTAACGAGTCACATC-FAM | 46 |
| 2a | 10-6 nt 5'-3' overhang | CGCGCGCATGTGACTCGTTACCTGAGTTTTTACTCAGGTAACGAGTCACATGATATATATAT | 62 |
| 2b | 10-6 nt 5'-3' overhang FAM | FAM-CGCGCGCATGTGACTCGTTACCTGAGTTTTTACTCAGGTAACGAGTCACATGATATATATAT | 62 |
| 3a | 6 nt 5' overhang | CGCGCGCATGTGACTCGTTACCTGAGTTTTTACTCAGGTAACGAGTCACATG | 52 |
| 3b | 6 nt 5' overhang FAM | FAM-CGCGCGCATGTGACTCGTTACCTGAGTTTTTACTCAGGTAACGAGTCACATG | 52 |
| 4a | 10 nt 3' overhang | CATGTGACTCGTTACCTGAGTTTTTACTCAGGTAACGAGTCACATGATATATATAT | 56 |
| 4b | 10 nt 3' overhang FAM | FAM-CATGTGACTCGTTACCTGAGTTTTTACTCAGGTAACGAGTCACATGATATATATAT | 56 |
| 5a | 24 nt ssDNA | AGAGAGAGAGAGAGAGAGAGAGAG | 24 |
| 5b | 24 nt ssDNA 5' FAM | FAM-AGAGAGAGAGAGAGAGAGAGAGAG | 24 |
